# Pseudoautosomal gene *SHOX* exhibits sex-biased random monoallelic expression and contributes to sex difference in height

**DOI:** 10.1101/2021.08.23.457290

**Authors:** Atsushi Hattori, Atsuhito Seki, Naoto Inaba, Kazuhiko Nakabayashi, Kazue Takeda, Kuniko Tatsusmi, Yasuhiro Naiki, Akie Nakamura, Keisuke Ishiwata, Kenji Matsumoto, Michiyo Nasu, Kohji Okamura, Toshimi Michigami, Yuko Katoh-Fukui, Akihiro Umezawa, Tsutomu Ogata, Masayo Kagami, Maki Fukami

## Abstract

Adult men are, on average, ∼13 cm taller than adult women. Although previous studies have suggested a significant contribution of sex chromosomal genes to sexual dimorphism in height, all attempts to identify a male-specific growth gene have failed. In the present study, we analyzed transcripts from cartilage tissues, and found that the expression of *SHOX*, a growth-promoting gene in the pseudoautosomal region on the X and Y chromosomes, was lower in females than in males. DNA methylation analyses showed that *SHOX* has some characteristics of genes subjected to X chromosome inactivation (XCI). These findings indicate that sex difference in human height is mainly ascribed to incomplete spreading of XCI on a pseudoautosomal gene. More importantly, RT-PCR of fibroblast clones revealed XCI-independent random clonal monoallelic expression of *SHOX*. We presume that during eutherian evolution, *SHOX* translocated from an autosome to the proto-sex chromosome without losing the epigenetic memory of random clonal monoallelic expression and subsequently underwent partial XCI. This study provides a novel model of epigenetic gene regulation leading to phenotypic diversity in humans.

## Introduction

The mean adult height of men is ∼ 13 cm more than that of women (1, 2). This sexual dimorphism is primarily ascribed to the effects of sex hormones and sex chromosomal genes (1). Considering that individuals with 46,XY gonadal dysgenesis are in general ∼8 cm taller than those with 46,XX gonadal dysgenesis (1), sex chromosomal genes likely play a more prominent role in this phenotype than sex hormones. To date, however, all attempts to identify a male-specific growth gene have failed. Actually, no major growth-controlling gene has been identified on the human sex chromosomes, except for *SHOX*, which resides in the pseudoautosomal region 1 (PAR1) of the X and Y chromosomes (3, 4). *SHOX* encodes a regulator of chondrocyte development, and its haploinsufficiency typically causes a height reduction by more than 10 cm (4, 5). *SHOX* was assumed to exert equivalent beneficial effects on linear growth of men and women, because all PAR1 genes including *SHOX* were believed to escape X chromosome inactivation (XCI) and show biallelic expression in both 46,XX and 46,XY cells (3, 6).

Here, we tested a new hypothesis that male-dominant expression of *SHOX* in the cartilage tissues contributes to the sex difference in height. This hypothesis is primarily based on the recent report by Tukiainen et al. that average transcript levels of PAR1 genes in various female tissues were lower than those in male tissues (7). The authors proposed that male-dominant expression of PAR1 genes reflects incomplete XCI, because (i) transcript levels of these genes from the inactive X chromosome (Xi) accounted for only ∼80% of those from the active X chromosome (Xa), and (ii) no systematic up- or down-regulation of PAR1 genes was observed on the Y chromosome (Y). However, Tukiainen et al. did not analyze transcripts in the cartilage tissues. Moreover, the authors did not investigate DNA methylation profiles, although it is known that XCI usually alters DNA methylation profiles of target genes (8).

## Results

### Transcriptome analysis for X chromosomal genes

We first examined sex differences in the transcript levels of X chromosomal genes. Microarray-based transcriptome analysis was carried out using cartilage tissues obtained from four adults (two women and two men) and cultured chondrocytes established from 12 children (six girls and six boys) (Table S1). The results revealed male-dominant expression of most PAR1 genes, along with female-dominant or sex-unbiased expression of known XCI-escape genes in the X-differential region (Fig. 1A). These results are consistent with previous data obtained from other human tissues (7). However, since *SHOX* expression remained low in most samples, we could not evaluate the difference in *SHOX* transcript levels between male and female samples.

**Fig. 1.**
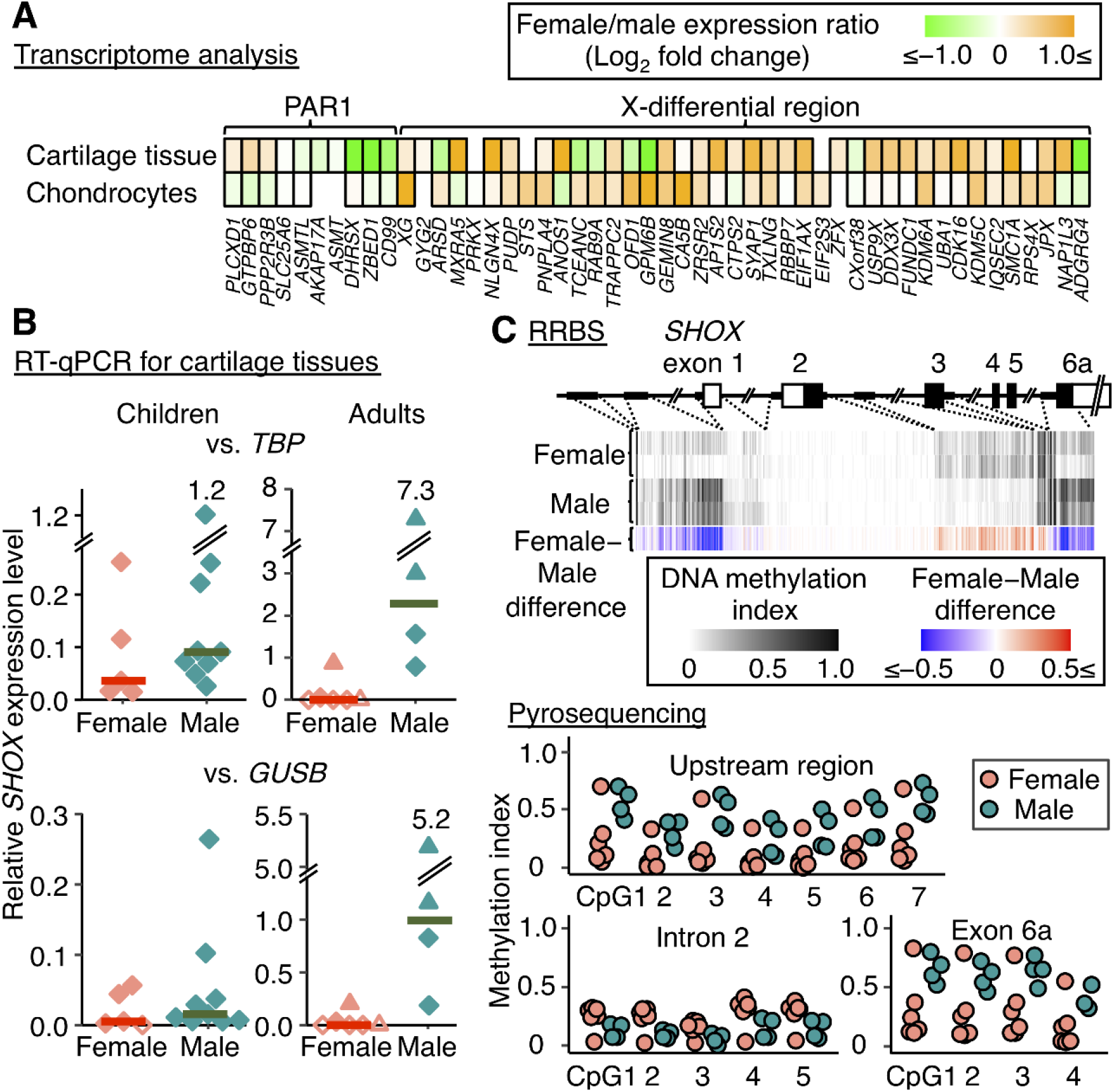
Representative results of mRNA quantification and DNA methylation analyses. **A**. Microarray-based transcriptome analysis for X chromosomal genes in cartilage tissues of women (n = 2) and men (n = 2) and cultured chondrocytes of girls (n = 2) and boys (n = 2). Female/male expression ratios of X chromosome inactivation (XCI)-escape genes are shown. PAR1, the short arm pseudoautosomal region. **B**. RT-qPCR for cartilage tissues of women (n = 6) and men (n = 4) and of girls (n = 5) and boys (n = 9). Relative *SHOX* expression levels against *TBP* and *GUSB* are shown. The diamond and triangle symbols indicate fresh frozen and RNAlater-treated tissues, respectively. The horizontal bars depict the median values of female and male samples. Unfilled symbols indicate values lower than the detection limit. **C**. DNA methylation analyses of the *SHOX*-flanking region. Upper panel, the results of reduced representation bisulfite sequencing (RRBS) for cartilage tissues of women (n = 2) and men (n = 2). The white and black boxes depict non-coding and coding regions of *SHOX*, respectively. For each CpG site, we calculated the difference in the mean methylation index between female and male samples (Female-male difference). Lower panel, the results of pyrosequencing for cartilage tissues of women (n = 6) and men (n = 4).

### RT-quantitative PCR (RT-qPCR) for *SHOX*

We examined *SHOX* expression levels in cartilage tissues of 10 adults (six women and four men) and 14 children (five girls and nine boys) using RT-qPCR. A transcript corresponding to *SHOX* exons 5 and 6a was quantified using TaqMan assays. Repeated experiments using two internal controls (*TBP* and *GUSB*) yielded consistent results. In both the adult and child groups, the median values of relative *SHOX* expression were higher in male samples than in female samples (Fig. 1B).

### DNA methylation analyses for X chromosomal genes

Next, to examine sex-specific DNA methylation of X chromosomal genes, we performed reduced representation bisulfite sequencing (RRBS) for cartilage tissues of four adults (two women and two men). RRBS is a method to assess DNA methylation profiles of CpG-rich genomic regions (9). The results showed that CpG sites in the X-differential region tended to be more highly methylated in female samples than in male samples, while most CpG sites in PAR1 showed similar methylation profiles in samples of both sexes (Fig. S1). These results can be explained by the presence of several XCI-subject genes only in the X-differential region (8), because such genes are characterized by relative hypermethylation in the regulatory regions (Fig. S2). However, we observed significant sex differences in the methylation indexes of three CpG clusters in the *SHOX*-flanking region (Fig. 1C and Fig. S1). Specifically, there were male-dominant DNA methylation of a 3.2 kb region which is 3.9 kb upstream of *SHOX* exon 1 (chrX: 578,055–581,224; GRCh37/hg19) and a 1.9 kb region in intron 5-exon 6a (chrX:604,185– 606,039). Furthermore, female-dominant DNA methylation was detected in a 1.2 kb region in intron 2 (chrX:592,562–593,801). Notably, the 1.2 kb intronic region contained a CpG island and enhancer/promoter-associated histone marks (H3K4Me1) (Fig. S3), indicating that this region may be involved in gene regulation.

The results of RRBS were validated by pyrosequencing for cartilage tissues of 10 adults (six women and four men) (Table S2). The results confirmed sex differences in the DNA methylation indexes of 16 CpG sites around *SHOX*, although there were some inter-individual variations (Fig. 1C). Our data imply that *SHOX* in female cartilage tissues is characterized by relative hypermethylation of the regulatory region and relative hypomethylation of the gene body. These methylation profiles likely reflect the condition on Xi, because Xa in women and the two sex chromosomes in men are all free from XCI and expected to share similar methylation profiles for PAR1 (10).

### Assessment of *SHOX* allelic expression in fibroblast clones

The aforementioned results indicate that *SHOX* expression in female cartilage tissues is partially suppressed possibly reflecting partial XCI. We attempted to confirm reduced *SHOX* expression from Xi. To this end, we examined *SHOX* allelic expression in 135 fibroblast clones established from single cells of five children (two girls and three boys). Chondrocytes were not used, because they tended to lose *SHOX* expression during cell culture. Unexpectedly, RT-PCR revealed monoallelic expression (MAE) of *SHOX* in several clones, along with biallelic or null expression in the remaining clones (Table 1, Fig. 2, and Table S3). *SHOX*-MAE in the clones was complete; in these clones, RT-PCR products of one allele were not detectable. Serial RT-PCR for multiple clones confirmed stable inheritance of *SHOX*-MAE, except for a few clones in which *SHOX* expression levels gradually decreased during cell culture. We confirmed that the *SHOX*-MAE clones carried no chromosomal deletions or amplifications that may affect *SHOX* allelic expression. *SHOX*-MAE in the clones cannot be ascribed to genomic imprinting or mutations in the regulatory/coding regions, because in each individual, both the maternally and paternally derived *SHOX* alleles were implicated in MAE. More importantly, *SHOX*-MAE was completely independent of XCI; in female clones with *SHOX*-MAE, *SHOX* was expressed from Xa and Xi with a similar frequency (Table 1). Likewise, in male MAE clones, *SHOX* was stochastically expressed from X or Y (Table 1). These results provide evidence that *SHOX* undergoes XCI-independent random clonal MAE in both male and female cells.

**Table 1.** *SHOX* allelic expression in fibroblast clones.

| Donor ID | Sex | Number of tested clones | Numbers of clones with each *SHOX* expression pattern |  |  |  |  |
| --- | --- | --- | --- | --- | --- | --- | --- |
|  |  |  | Biallelic | MAE (Xa) | MAE (Xi) | MAE (Y) | Null |
| CS1 | F | 50 | 2 (4%) | 6 (12%) | 5 (10%) | ... | 37 (74%) |
| CS2 | F | 24 | 3 (13%) | 1 (4%) | 2 (8%) | ... | 18 (75%) |
| CS3 | M | 32 | 0 (0%) | 3 (9%) | ... | 2 (6%) | 27 (84%) |
| CS4 | M | 17 | 2 (12%) | 2 (12%) | ... | 2 (12%) | 11 (65%) |
| CS5 | M | 12 | 0 (0%) | 1 (8%) | ... | 0 (0%) | 11 (92%) |

MAE, monoallelic expression; Xa, active X chromosome; Xi, inactive X chromosome; Y, Y chromosome; F, female; M, male.

**Fig. 2.**
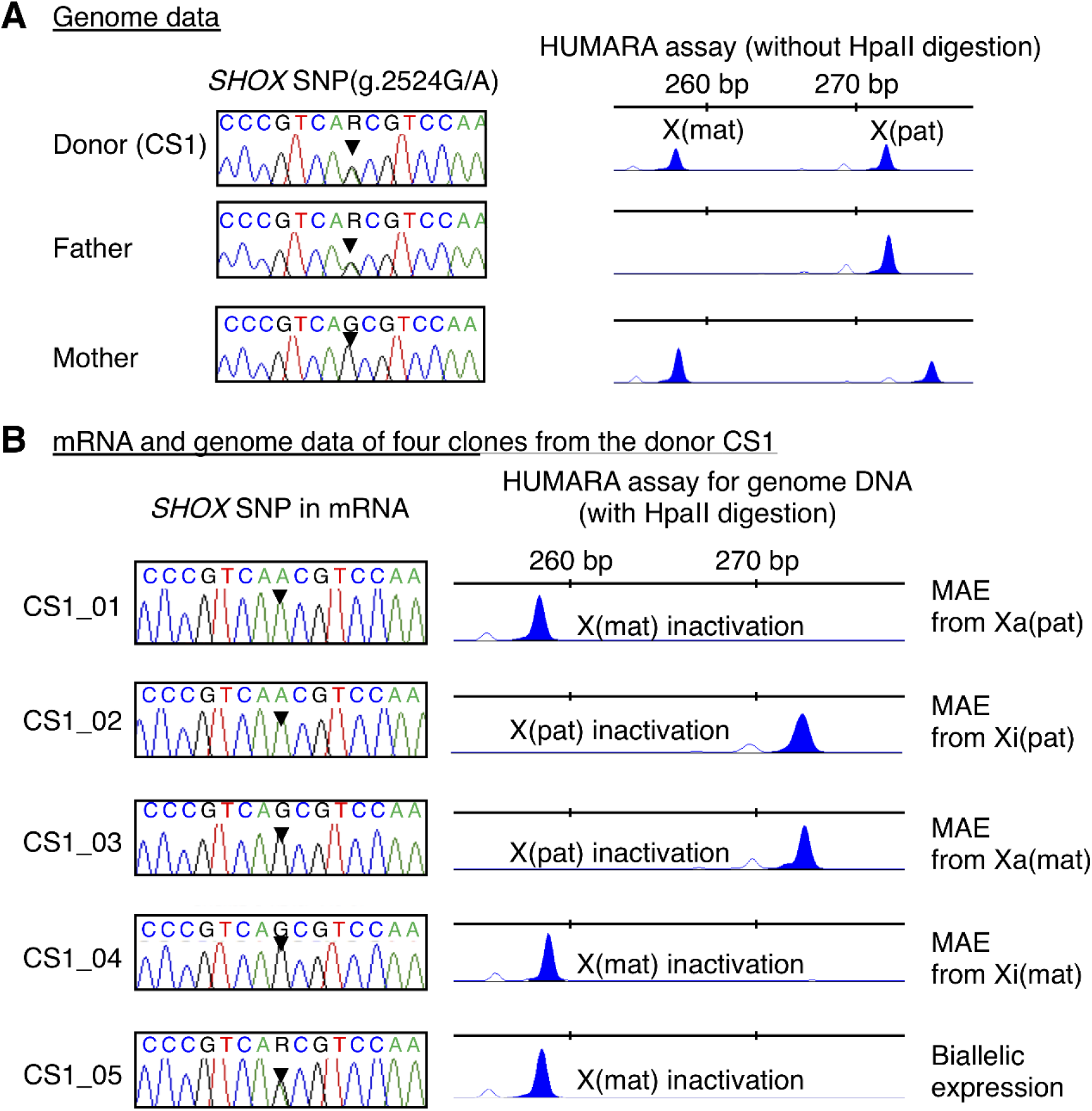
Representative results of *SHOX* allelic expression analysis in fibroblast clones. **A**. Genome data of a girl [CS1] and her parents. Left panels show the sequences around the informative *SHOX*-SNP. Arrowheads depict the polymorphic nucleotide. Right panels show the results of HUMARA assays. The 258 bp and 273 bp peaks in the girl correspond to the X chromosomes inherited from her mother [X(mat)] and father [X(pat)], respectively. **B**. mRNA and genome data of fibroblast clones with different types of *SHOX* allelic expression. In clones with *SHOX* monoallelic expression (MAE), *SHOX* was expressed from the active X(pat) and X(mat) [Xa(pat) and Xa(mat)], as well as from the inactive X(pat) and X(mat) [Xi(pat) and Xi(mat)].

### RT-qPCR and DNA methylation analyses for clones

We performed RT-qPCR for fibroblast clones, with the aim of clarifying whether *SHOX* transcripts from Xi are fewer than those from Xa or Y. However, because biallelic *SHOX* expression was observed only in a few clones (Table 1), and *SHOX* expression was low in most fibroblast clones (Fig. S4A), it was impossible to compare *SHOX* expression levels from Xi, Xa, and Y.

We also examined whether *SHOX* allelic expression was associated with specific DNA methylation profiles of the CpG sites around *SHOX*. Pyrosequencing for 12 *SHOX*-flanking CpG sites revealed that DNA methylation indexes of clones were highly variable and were not correlated with *SHOX* expression patterns (Fig. S4B).

### Whole exome sequencing for genomic DNA of a female donor and mRNA of her clones

Lastly, we investigated whether any X chromosomal genes other than *SHOX* show random clonal MAE. We performed whole exome sequencing for genomic DNA obtained from one female donor (sample ID, CS1) and mRNA extracted her four clones. These clones exhibited different patterns of *SHOX*-MAE (MAE from paternally derived Xa, maternally derived Xa, paternally derived Xi, and maternally derived Xi). As a result, we identified 38 X chromosomal genes that showed MAE or skewed biallelic expression in three or more clones (Table S4).

However, all of these MAE cases were likely to reflect XCI, because the expressing alleles co-segregated with the XCI pattern of the clones. Moreover, 32 of the 38 genes have previously been described as XCI-subject or -mostly subject genes (11). Although the remaining six genes have been reported as XCI-escape genes (11), Xa-specific expression of these genes is indicative of XCI. Altogether, this experiment provided no evidence that an X chromosomal gene other than *SHOX* shows random clonal MAE.

## Discussion

This study showed that *SHOX* expression levels were lower in cartilage tissues of females than in those of males. Since *SHOX* is known to facilitate linear growth in a dosage-sensitive manner (4), relatively low *SHOX* expression in female cartilage tissues likely contributes to short stature of women. This notion is supported by the fact that female patients with *SHOX* haploinsufficiency usually manifest more severe phenotypes than male patients (4, 5). Moreover, similar phenotypes of male patients with *SHOX* mutations on X and Y (Table S5) affirm that *SHOX* is equally expressed from the two sex chromosomes in men. The lack of *SHOX* orthologs in rodents and several other species (4) may explain why sex-biased expression of *Shox* has not been reported in animals.

We observed sex-biased DNA methylation in three CpG clusters around *SHOX*. Previous studies have shown that in general, DNA methylation profiles were similar between Xa of women and the single X of men (10), and between XCI-escape and -subject genes on Xa (8). In contrast, on Xi, the average methylation rates of the promoter/enhancer regions of XCI-subject and -escape genes were reported to be 70% and 11% respectively, whereas those of the gene bodies were 64% and 75% respectively (8). The present study detected male-dominant DNA methylation of CpG sites around *SHOX* exons 1 and 6a, and female-dominant methylation of CpG sites in intron 2. Thus, *SHOX* has some characteristics of XCI-subject genes, i.e., relative hypermethylation of the regulatory region and relative hypomethylation in the gene body (8). This indicates partial XCI on *SHOX*. Consistent with this, Sun et al. reported diverse phenotypes of two women with the same *SHOX* deletion, who had extremely skewed XCI (12).

The next question was whether *SHOX* expression levels from Xi are lower than that from Xa and Y. Thus, we analyzed *SHOX* allelic expression in fibroblast clones established from single cells of five children. Unexpectedly, we observed *SHOX*-MAE in several clones. *SHOX*-MAE in these clones was mitotically stable and completely independent of the parental-origin or XCI. These results provide the first evidence that a sex chromosomal gene can undergo random clonal MAE. It was impossible to compare *SHOX* expression levels from Xi, Xa, and Y, because biallelic *SHOX* expression was observed only in a few clones and *SHOX* transcript levels were low in most clones.

Random clonal MAE is defined as stochastic silencing of one allele in a cell, which is stably transmitted to all daughter cells (13–18). Random clonal MAE is regarded as a biologically important phenomenon that produces cellular heterogeneity in a tissue (13–18). Yet, only a few autosomal genes have been implicated in this phenomenon (13, 15). Actually, although single-cell assays have detected MAE of several genes, most of the cases were explicable by mutations in the regulatory/coding regions, imprinting, or low transcriptional levels (13, 18–20). Moreover, frequent MAE of X chromosomal genes was invariably attributed to XCI (6, 13). Our data challenge the current concept of random clonal MAE. Notably, in MAE clones, *SHOX* was expressed from Xa and Xi with similar frequencies. This indicates that all processes of XCI, including XIST-induced chromatin condensation, do not affect MAE. Moreover, *SHOX* displayed not only MAE, but also biallelic and null expression. Such variations have also been reported for autosomal MAE genes (13). This suggests that “random clonal MAE” does not represent simple silencing of one allele. Instead, this phenomenon probably specifies a unique group of genes, whose allelic expression is randomly turned on or off during early development, and once the decision is made, the information is stably maintained across cell divisions. Our findings imply that human skin tissues comprise heterogeneous cell populations with different types of *SHOX* allelic expression (Fig. 3). If this is also applicable to cartilage tissues, such cellular diversity may contribute to inter-individual height variations. On the other hand, since pyrosequencing for fibroblast clones detected no association between *SHOX* allelic expression and DNA methylation, the regulatory mechanism of *SHOX*-MAE remains to be clarified. Indeed, previous studies have shown that random clonal MAE of autosomal genes does not necessarily entail DNA methylation changes (13).

**Fig. 3.**
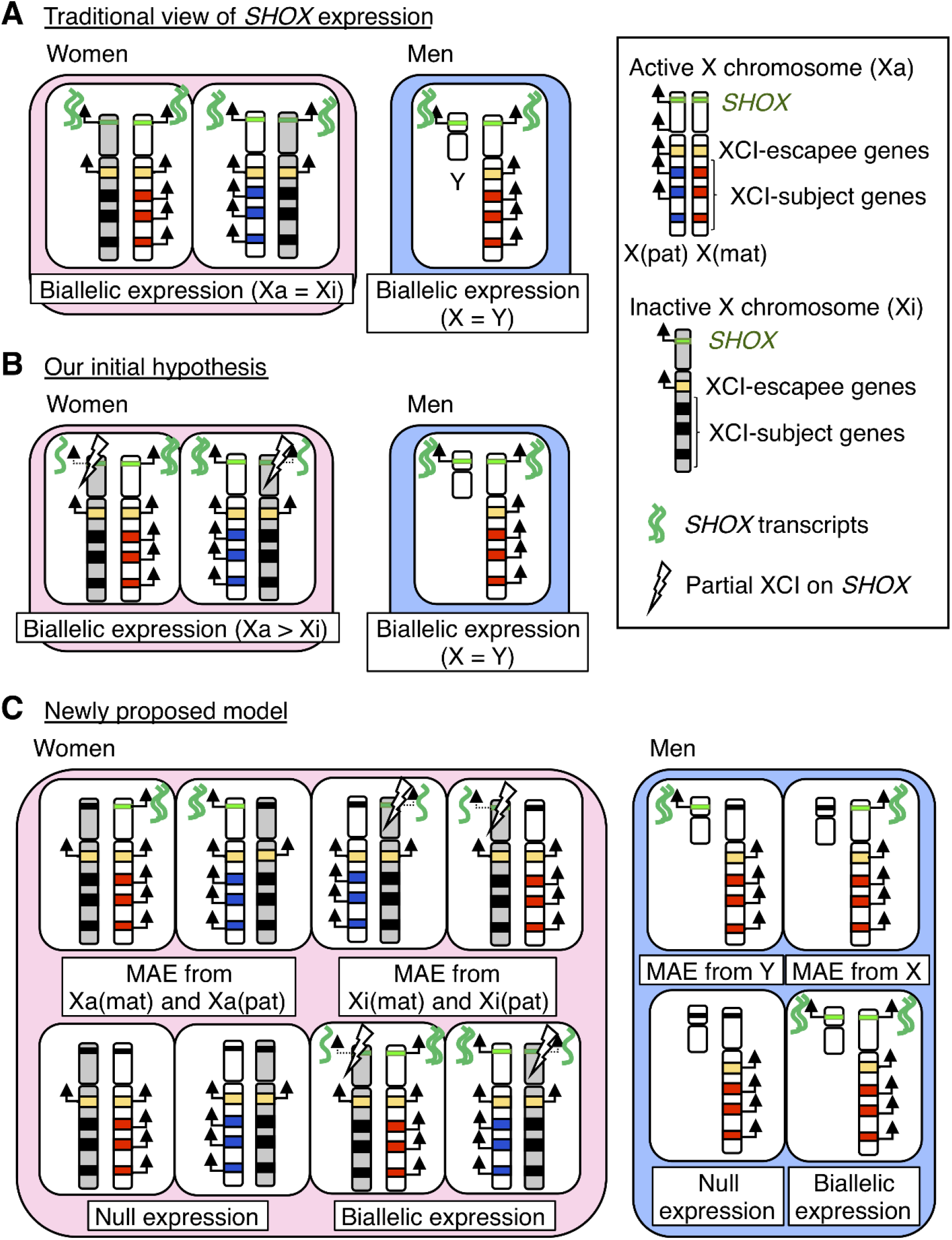
Predicted models of *SHOX* allelic expression. **A**. Traditional view of *SHOX* expression. *SHOX* was believed to be equally expressed from active X (Xa) and inactive X (Xi) in women and from X and Y in men. X(pat) and X(mat) indicate X chromosomes derived from the father and mother, respectively. **B**. Our initial hypothesis. We speculated that *SHOX* exhibits biallelic expression, but its expression from Xi is lower than that from Xa. **C**. A newly proposed model. *SHOX* exhibits random monoallelic expression (MAE), as well as biallelic and null expression. *SHOX*-MAE is independent of X chromosome inactivation (XCI). Thus, cartilage tissues likely consist of heterogeneous cell populations with different types of *SHOX* allelic expression. *SHOX* expression from Xi appears to be reduced due to partial XCI.

Lastly, we examined whether other X chromosomal genes also show random clonal MAE. However, whole-exome sequencing identified no additional genes with random clonal MAE. These findings are consistent with the report by Garieri et al., which demonstrated extremely skewed allelic expression of *SHOX*, but not that of other PAR1 genes (21). Since random clonal MAE of sex chromosomal genes has not been documented in any animal species, *SHOX* may be an exceptional gene on the sex chromosomes. Of note, *SHOX* has a unique evolutionary history. During eutherian evolution, PAR1 including *SHOX* translocated from an autosome to the proto-sex chromosome (22). We speculate that *SHOX* emerged as a random clonal MAE gene on an autosome, then translocated to the proto-sex chromosome without losing the memory of random MAE, and was finally subjected to partial XCI. In this scenario, the other PAR1 genes probably maintained biallelic expression throughout evolution. However, we cannot exclude the possibility that other sex chromosomal genes exhibit random clonal MAE, because our whole-exome analyses focused only on genes that were strongly expressed in fibroblasts and contained informative SNPs in the tested DNA sample.

In conclusion, the results of this study indicate that sex differences in height are mainly ascribed to partial XCI on a pseudoautosomal gene, rather than the presence of male-specific growth genes. Furthermore, our data imply that *SHOX* is an exceptional sex chromosomal gene that retains an epigenetic memory of autosomal random clonal MAE. This study provides novel insights into the transcriptional regulation of human genes generating phenotypic diversity.

## Materials and Methods

### The target gene *SHOX*

In this study, we focused on *SHOX*a, the major transcript of *SHOX* (NM_000451.3). *SHOX*a consists of exons 1–5 and 6a and resides at chrX:585,079–607,558 and chrY: 535,079–557,558 (GRCh37/hg19). Although previous studies have identified several splice variants of *SHOX*, all of these variants except for *SHOX*a are of unknown clinical importance (23).

### Human samples

The human samples analyzed in this study are summarized in Supplementary Table S1. The samples were obtained from unrelated individuals. First, we analyzed postmortem cartilage tissues obtained from the knee joints of 10 adults (six women and four men). These samples were purchased from Articular Engineering (Catalog IDs, CDD-H-6000-N-1G-R and CDD-H-6000-N-1G-F; Northbrook, IL, USA). The donors of these samples died of trauma or non-endocrine disorders (Table S1). The samples were either frozen (n = 6) or stored in the RNAlater reagent (Thermo Fisher Scientific, Waltham, MA, USA) (n = 4).

Second, we examined cartilage tissues and cultured chondrocytes obtained from 26 healthy children. These samples were taken in our hospital during surgery for polydactyly. Of these, 14 cartilage tissues from five girls and nine boys were frozen in liquid nitrogen immediately after surgery. The remaining 12 cartilage tissues from six girls and six boys were subjected to chondrocyte isolation. These tissues were pulverized and cultured in Dulbecco’s modified Eagle’s medium containing 17% fetal bovine serum. We confirmed that the cells had chondrocyte-compatible morphological characteristics (24, 25) and expressed *COL2A1*, a marker of chondrocytes.

Lastly, we analyzed skin fibroblast clones established from single cells of surgical specimens. These skin samples were obtained from two girls and three boys with polydactyly. To determine the parental origin of the X chromosomes in the clones, we collected saliva samples from the donors’ parents using the Oragene-DNA kit (DNA Genotek, Ontario, Canada).

### Primer information

Primers used in this study is listed in Table S2.

### Extraction of total RNA and genomic DNA

Total RNA of cartilage tissues was extracted using CRYO-PRESS (Microtec, Funabashi, Japan) and the TRIzol reagent (Thermo Fisher Scientific), and that of cultured cells was obtained using the AllPrep DNA/RNA/miRNA Universal kit or the miRNeasy mini kit (QIAGEN, Hilden, Germany). The samples were reverse-transcribed into cDNA with the High-Capacity cDNA reverse transcription kit (Thermo Fisher Scientific).

Genomic DNA of cartilage tissues and cultured cells was extracted using the Gentra Puregene kit (QIAGEN) or the AllPrep DNA/RNA/miRNA Universal kit (QIAGEN). Genomic DNA of saliva was extracted using the prepIT-L2P kit (DNA Genotek).

### Transcriptome analysis for X chromosomal genes

Transcriptome analyses were performed for cartilage tissues of adults (two women and two men) and cultured chondrocytes of children (six girls and six boys) using catalog one-color microarrays (SurePrint G3 Human Gene Expression microarray, 8 × 60 k format; Agilent Technologies, Santa Clara, CA, USA). The data were analyzed using GeneSpring software (version 14.9, Agilent Technologies). We focused on genes on the X chromosome, particularly those that escape XCI. Transcripts with low expression levels and low signal quality were filtered out. Specifically, we excluded probes that were assessed as “not uniform”, “saturated”, or “population outliers” in one or more of the tested samples. We also excluded probes that were “not significant” or “not above background” in one or more samples of both sex groups. The female-male ratio at each transcript level was calculated by subtracting the average value of logarithmically transformed signal intensities in female samples from that in male samples.

### RT-qPCR for *SHOX*

*SHOX* expression levels in the cartilage tissues of adults (six women and four men) and children (five girls and nine boys) were analyzed by RT-qPCR. The transcript corresponding to exons 5 and 6a of *SHOX* (assay ID, Hs00757861_m1; Thermo Fisher Scientific) was quantified using TaqMan assays on a 7500 Fast real-time PCR system (Thermo Fisher Scientific). *SHOX* expression levels were calculated using the ΔΔ-CT method against two internal control genes, *TBP* (assay ID, Hs00427620_m1) and *GUSB* (assay ID, Hs00939627_m1). Each sample was analyzed in triplicate.

### RRBS for X chromosomal genes

We performed RRBS for cartilage tissues of four adults (two women and two men). RRBS libraries were prepared according to the standard methods with some modifications (26, 27). In brief, genomic DNA samples were sonicated and treated with proteinase K (Thermo Fisher Scientific) and RNase A (Nacalai Tesque, Kyoto, Japan). Purified samples were digested with MspI (New England BioLabs, Ipswich, MA, USA), subjected to gap filling and A-tailing with the Klenow fragment (Thermo Fisher Scientific), and ligated with the NEBNext methylated adaptor (New England BioLabs). The adaptor-ligated DNA samples were subjected to bisulfite conversion by the EZ DNA Methylation-Gold kit (Zymo Research, Irvine, CA, USA) and amplified by PCR with the KAPA HiFi HotStart Uracil+ ReadyMix kit (Roche, Basel, Switzerland) (Table S2). The libraries were subjected to 150-bp paired end sequencing on a next-generation sequencer (NextSeq, Illumina, San Diego, CA, USA).

Sequence reads were mapped to the human reference genome for bisulfite sequencing (CT-converted hs37d5) using the Bismark program (28). Adaptor trimming and quality control were performed using Trim Galore software (http://www.bioinfor-matics.babraham.ac.uk/projects/trim_galore/). The methylation index of each CpG site was calculated using the methylKit package (v.1.10.0) in R (29). We excluded CpG sites whose read-depth was < 10 in any of the tested samples. We also excluded constantly hypermethylated CpG sites with methylation rates of > 80% in all samples. Sex differences in the DNA methylation levels were calculated by subtracting the mean methylation index of male samples from that of female samples.

### Pyrosequencing for *SHOX*-flanking CpG sites

We performed pyrosequencing for the *SHOX*-flanking CpG sites using cartilage tissues from 10 adults (six women and four men). Genomic DNA samples were treated with bisulfite using the EZ DNA Methylation-Gold kit (Zymo Research). Three genomic intervals encompassing CpG clusters (chrX:580,908–581,116, 593,482–593,694, and 605,642–605,771) were PCR-amplified and subjected to pyrosequencing on PyroMark Q24 (QIAGEN) (Table S2). We analyzed the methylation indexes of 16 CpG sites (seven in the *SHOX* upstream region, five in intron 2, and four in intron 5-exon 6a).

### *In silico* analysis for the region of female-dominant methylation in *SHOX* intron 2

We performed in silico analyses for the 1.2 Kb region in *SHOX* intron 2, which showed female-dominant DNA methylation. We referred to the UCSC genome browser (https://genome.ucsc.edu) to examine whether the region contains CpG islands or promoter/enhancer-associated histone marks.

### Establishment of clones from single fibroblasts

Fibroblast clones were established from single cells of five children (three boys and two girls). We did not use chondrocytes, because we found that they frequently lost *SHOX* expression during cell culture. Fibroblast clones were established using the following procedure. First, we screened our banked surgical specimens by Sanger sequencing of *SHOX* exons. As a result, we identified six skin samples whose donors carried heterozygous exonic SNPs in *SHOX*. Then, we sequenced genomic DNA samples of the donor’s parents, to determine trio-genotypes. The results showed that five of the six samples had informative trio-genotypes, i.e., one of the donor’s parents was homozygous for one allele of the *SHOX*-SNP, and the other parent was homozygous or heterozygous for the alternative allele. Thus, single cells were isolated from the five skin samples by using a single-cell picking system (ASONE, Osaka, Japan). The cells were grown in DMEM containing 17% FBS and antibiotics to reach ∼90% confluence in 6-well or 10-cm plates. Genomic DNA and total RNA were extracted from the cells, as described above.

### Assessment of *SHOX* allelic expression in clones

First, we screened *SHOX*-expressing clones by RT-PCR using primers on *SHOX* exons 2 and 3 (Table S2). For each *SHOX*-expressing clone, we amplified and sequenced a cDNA fragment encompassing the informative *SHOX*-SNP of the donor (Table S2). We calculated the ratio of the reference allele transcript to total transcripts (“the reference allele ratio”) from the amplitudes of base call peaks. The sangerseqR package in R (30) was used to calculate the ratio. Consequently, the clones were classified into four groups according to the expression patterns of *SHOX*; (i) biallelic expression (reference allele ratios of between 0.2 and 0.8), (ii) skewed biallelic expression (reference allele ratios of between 0.1 and 0.2 or between 0.8 and 0.9), (iii) MAE (reference allele ratios of ≤ 0.1 or ≥ 0.9), and (iv) null expression (31). For clones with *SHOX* null expression, we amplified *COL1A1* mRNA to confirm the quality of the mRNA sample. Stable maintenance of *SHOX* expression patterns across mitotic cell divisions was confirmed by serial RT-PCR of multiple clones.

The parental origin of the *SHOX*-expressing allele(s) in each clone was determined from the trio-genotype. Furthermore, for clones from female donors, genomic DNA samples were subjected to methylation-specific microsatellite assays of the androgen receptor gene (“the HUMARA assay”), with the aim of determining the parental origin of Xa and Xi. The methods of the HUMARA assay were described previously (32).

### Chromosomal copy-number analysis of clones

All *SHOX*-expressing clones were subjected to copy-number analysis, because cultured cells are known to have a risk of acquiring new chromosomal abnormalities during mitosis (14). Genomic DNA from each clone was analyzed by microarray-based comparative genomic hybridization using catalog whole-genome microarrays (SurePrint G3 Human CGH microarray, 8 × 60 k format, Agilent technologies) and/or by multiplex ligation dependent probe amplification using the *SHOX* SALSA MLPA Probemix (assay ID, P018-G2; MRC Holland, Amsterdam, The Netherland). Clones with apparent copy-number abnormalities were excluded from further analyses.

### RT-qPCR and DNA methylation analyses for clones

RT-qPCR was performed for total RNA extracted from eight fibroblast clones. Furthermore, we performed pyrosequencing for genomic DNA samples of seven clones with biallelic, monoallelic, and null expression of *SHOX*. A total of 12 CpG sites in the *SHOX*-flanking region were analyzed.

### Whole exome sequencing for genomic DNA of a female donor and mRNA of her clones

We performed whole exome sequencing of a genomic DNA sample from a female donor (sample ID, CS1) and mRNA samples of her four clones. In these clones, *SHOX* was expressed from different X chromosomes, i.e., paternally derived Xa, maternally derived Xa, paternally derived Xi, and maternally derived Xi. Sequencing was performed by Macrogen Japan (Kyoto, Japan). We called exonic variants that were heterozygous in the genome of the donor. Variants with low quality scores and those with low expression levels (read depth ≤ 10 in all clones) were excluded. Then, we calculated the reference allele ratio of each variant in mRNA samples and searched for genes with MAE (reference allele ratios of ≤ 0.1 or ≥ 0.9) or skewed allelic expression (reference allele ratios of between 0.1 and 0.2 or between 0.8 and 0.9). The reference allele ratios were calculated from read depths of mRNA sequencing. Lastly, we examined whether MAE of the genes could be ascribed to XCI. To this end, we investigated whether the gene was expressed exclusively from Xa, and whether the gene has been described as an XCI-subject gene in the literature (11).

## Acknowledgments

This study is funded by Japan Society for the Promotion of Science 17H06428 (to M.F.), Japan Agency for Medical Research and Development 20ek0109464h0001 (to M.F.), National Center for Child Health and Development 2019A-1 (to M.F.), and The Takeda Science Foundation (to M.F.)

## Competing Interest Statement

The authors declare no competing interest.

## Notes

### Competing Interest Statement

The authors have declared no competing interest.

