## Supplementary Figures for "Pseudoautosomal gene *SHOX* exhibits sex-biased random monoallelic expression and contributes to sex difference in height"

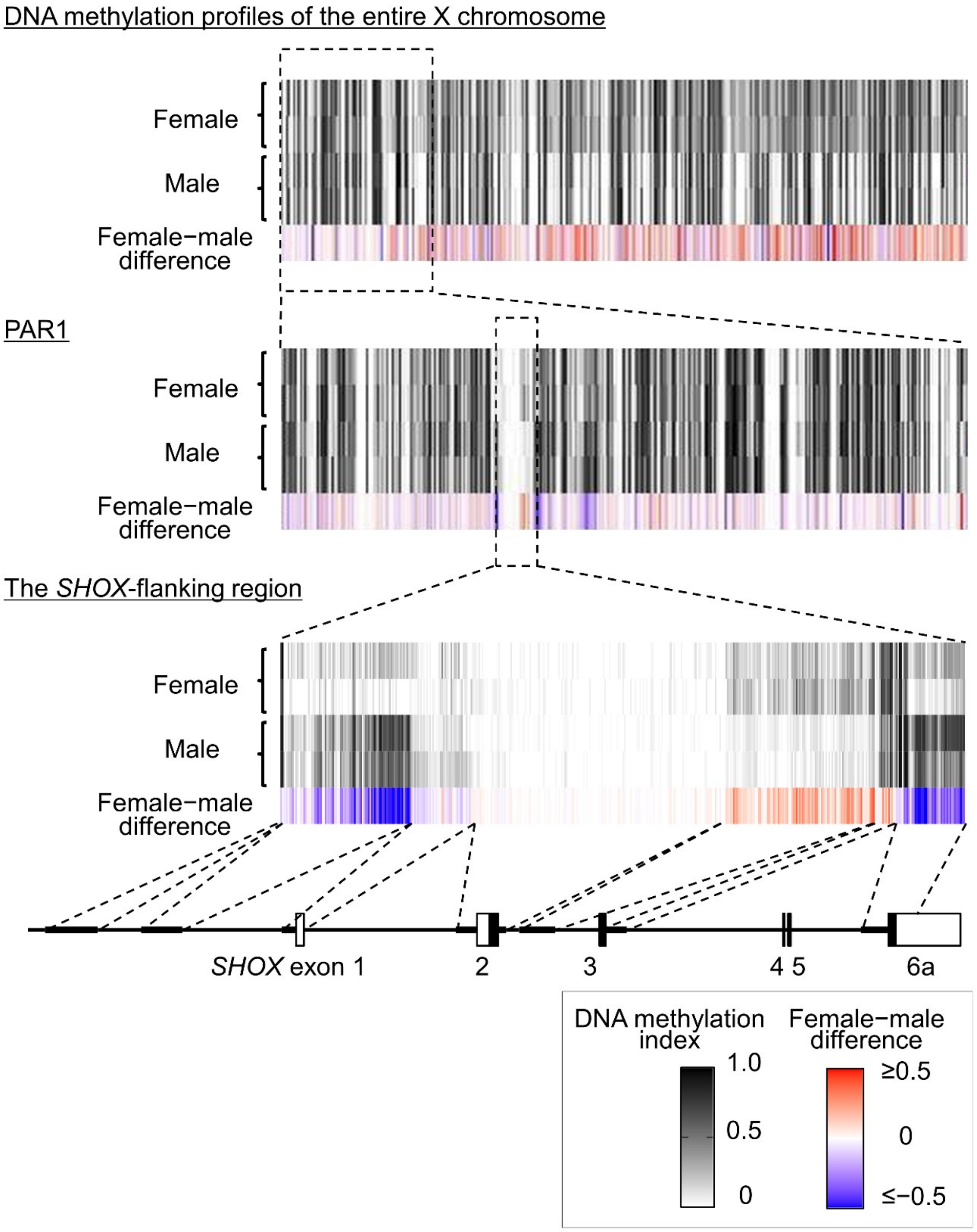


**Fig. S1. Results of reduced representation bisulfite sequencing (RRBS) for cartilage tissues.**

For each CpG site, we calculated the Female-Male difference (difference in the median methylation index between female and male samples). The white and black boxes depict non-coding and coding regions of *SHOX*, respectively. PAR1, pseudoautosomal region 1.

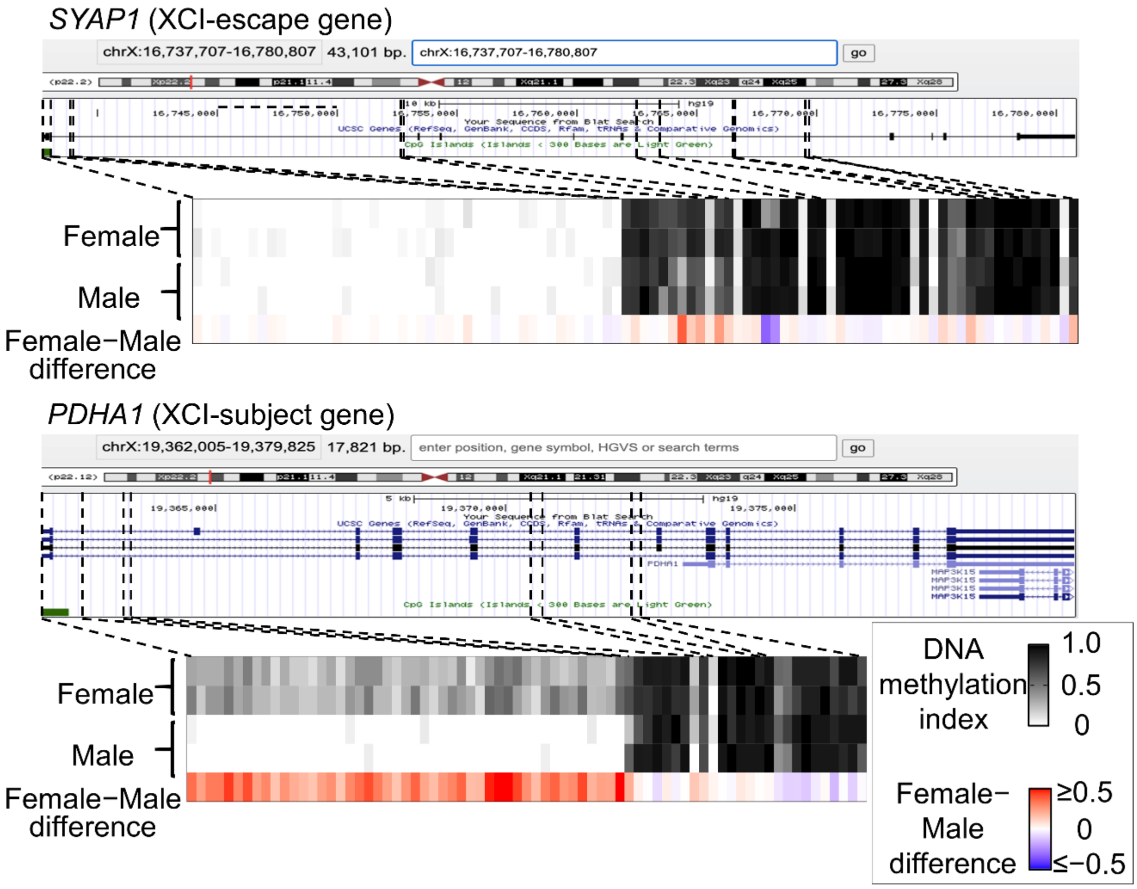


**Fig. S2. Representative results of reduced representation bisulfite sequencing (RRBS) of X chromosome inactivation (XCI)-escape and -subject genes.**

DNA methylation profiles of *SYAP* (an XCI-escape gene) and *PDHA1* (an XCI-subject gene) are shown. For each CpG site, we calculated the Female-Male difference (difference in the median methylation index between female and male samples).

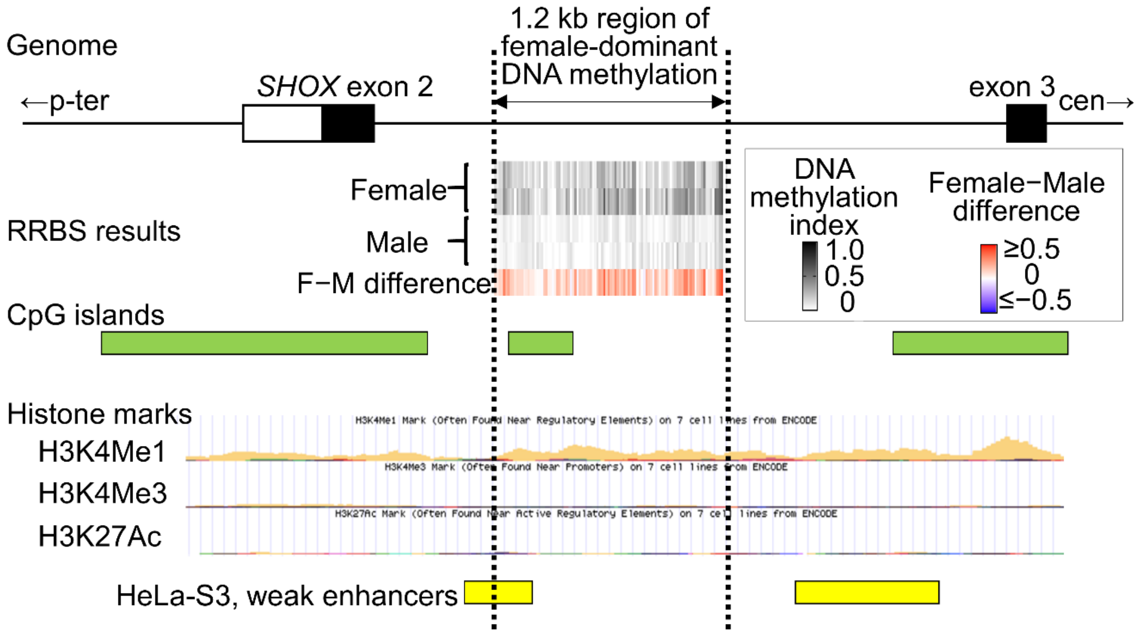


**Fig. S3. Representative *in silico* analysis for the region of female-dominant DNA methylation in *SHOX* intron 2.**
White and black boxes indicate non-coding and coding regions of *SHOX*, respectively. The 1.2 kb region contained a CpG island and was associated with H3K4Me1 in H1-hESC and a weak enhancer in HeLa-S3.

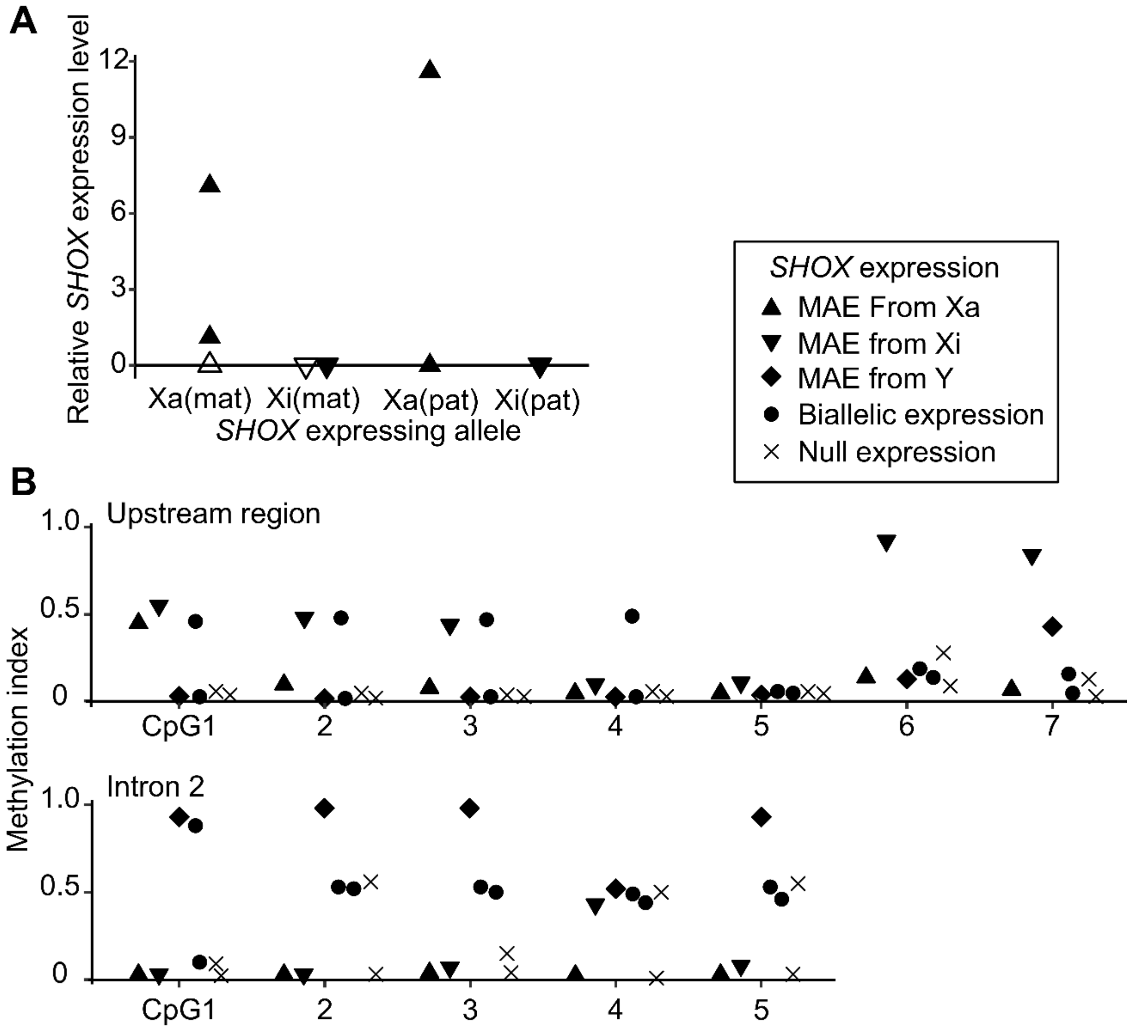


**Fig. S4. *SHOX* RT-qPCR and pyrosequencing for fibroblast clones.**

**A.** Relative expression levels of *SHOX* against *TBP*. The results of fibroblast clones expressing *SHOX* from maternally derived active X [Xa(mat)], maternally derived inactive X [Xi(mat)], paternally derived active X [Xa(pat)], and paternally derived inactive X [Xi(pat)] are shown. Each symbol represents a single clone. Open triangles indicate values lower than the detection limit.

**B**. DNA methylation profiles of the upstream region and intron 2 of *SHOX*. The results of clones with different types of *SHOX* allelic expression are shown. Each symbol represents a single clone. MAE, monoallelic expression; Xa, active X; Xi, inactive X.
